# In Silico Screening and Analysis of SNPs in Human ABCB1 (MDR1) Gene

**DOI:** 10.1101/505859

**Authors:** Amira Hussien, Alaa A.M. Osman

**Affiliations:** Department of Pharmacology, Faculty of Pharmacy, University of Gezira, Sudan; Department of Clinical Pharmacy and Pharmacy Practice, Faculty of Pharmacy, University of Gezira, Sudan

## Introduction

ABCB1 is one of the integral membrane adenosine triphosphate (ATP)-binding cassette (ABC) transporter proteins superfamily. To date, the human ABC transporter superfamily contains 49 members that classified into seven subfamilies (ABCA to ABCG). These transport proteins are localized to various membranes of cellular organelles (except for ABCE and ABCF), where they function as ATP dependent, and unidirectional transmembrane efflux pumps to various endogenous and exogenous compounds (1–7). Being efflux pumps, these transporters exert a wide variety of physiological roles including protective, excretory and regulatory functions. They create barriers between systemic blood circulation and many organs such as brain, cerebral spinal fluid, placenta and testis, thus restrict permeability of drugs and toxic compounds and hence protect these organs. They also expressed in the liver and kidney, where they excrete xenobiotics and endogenous compounds. In addition, they limit absorption of drugs into the systemic circulation by effluxing drugs into the gastrointestinal tract. Furthermore, ABCB transporters regulate many endogenous molecules affecting lipid and bile acid synthesis, antigen presentation, heme and iron homeostasis, steroid hormones transport and homeostasis, and signaling molecules such as cyclic nucleotides and ions, etc (1–8).

ABCB1 was formerly termed P-gp (Permeability-glycoprotein) as it thoughts to play a role in modulating cellular permeability to drugs (7). Thereafter many studies confirmed that P-gp is a transporter glycoprotein that has a major role in drug effluxes and resistance to several structurally and biochemically unrelated drugs including anticancer (e.g., paclitaxel, irinotecan), antibiotics (e.g., erythromycin, levofloxacin), immunosuppressants (e.g., ciclosporin, tacrolimus), cardiac drugs (e.g., digoxin, quinidine), calcium channel antagonists (e.g., diltiazem, verapamil), HIV protease inhibitors (e.g., ritonavir, saquinavir), corticosteroids, lipid-lowering agents, B-Adrenoceptor blockers, and H1 and H2 antihistamines, among others; hence termed multidrug resistant (MDR) (1–8).

Humans ABCB1 gene is located on chromosome 7q21, 25 kb from ABCB4 with a length ~100 kb including 32 exons (9). This gene encodes ABCB1 protein which is a 170-kD transmembrane glycoprotein of an approximate length of 1280 amino acids residues. ABCB1 composed of two halves, each consist of six hydrophobic transmembrane α-helices and an ATP binding domain (1–3, 5–8).

More than fifty single nucleotide polymorphisms (SNPs) in the ABCB1 protein coding sequence were identified (2, 4, 8). Of these, three SNPs are the most widely investigated: the synonymous SNPs rs1045642 (C3435T, Ile1145Ile) located on exon 26 and rs1128503 (C1236T, Gly412Gly) located on exons 12, and the non-synonymous (ns) SNP rs2032582 (G2677A/T, Ala893Ser/Thr) located on exon 21 (2–6, 8). The effects of these SNPs on the ABCB1 expression, disease susceptibility or its functions are subjected to controversial results in the literature (2–7).

Several studies reported that ABCB1 SNPs were associated with the risk of developing many diseases including cancer, ulcerative colitis, Parkinson disease, HIV-1, hypertension and rheumatoid arthritis among others (2, 5–7). Regarding cancer; a large meta-analysis of 39 independent studies found that the 3435T allele was associated with overall cancer risk (OR: 1.18, 95% CI: 1.04-1-34), although the risk is not high as compared with other cancer risks, but the risk is increased when stratified according to specific cancer subtypes (breast cancer, OR: 1. 42, 95% CI: 1.04-1.94; renal cancer, OR: 1.77 95% CI: 1.28-2.46; hematological malignancies, OR: 1.27, 95% CI: 1.10-1.46) (2, 10).

ABCB1 SNPs affect the pharmacokinetics and therapy outcomes (both side effects and clinical response) of many drugs especially antineoplastic agents (e.g., irinotecan, platinum, taxanes, anthracyclines, vinca alkaloids), antiretroviral therapy, antidepressants (e.g., fexofenadine, paroxetine), immunosuppressants (e.g., methotrexate, tacrolimus, cyclosporine, steroid) and digoxin, among others (2, 5–7, 11). The effects of SNPs are substrate dependent because different alleles have been associated with different drugs (2). Examples of effects on pharmacokinetics; the 3435T allele showed increased concentrations and decreased dose administration of tacrolimus (2, 12), also 3435T allele showed higher steady-state plasma concentrations of oral digoxin (5, 6, 13). Examples of effects on drug response; the 3435C allele showed increased platinum-based chemotherapy response in lung cancer (2, 14, 15). Another example, acute myeloid leukemia patients homozygous for the wild-type allele of ABCB1 at any locus (exons 12, 21 and 26) exhibited a significantly decreased overall survival with a higher probability of relapsed (16). Examples of effects on adverse effects; the C3435T mutation is associated with increased risk for nortriptyline-induced postural hypotension (17) and G2677T/A mutation predict tacrolimus neurotoxicity (18).

There is a huge number of SNPs known up to date, so it is not feasible to study all SNPs. Different techniques and tools are used to select the most damaging SNPs and to predict their effects on protein structure, stability and function. Structural bioinformatics is an area of bioinformatics focused on the structure, movement and interaction of biological macromolecules in 3-dimensional (3D) space. This approach is based on retrieving SNPs from databases and then filtering them using different tools. These tools usually fall into one of two categories. The first category is made up of tools that make predictions based solely on the protein sequence (e.g., SIFT, PROVEAN, PANTHER, PhD SNP), while the second one is made up of tools that incorporate structural information when making predictions (e.g., PolyPhen-2, SNAP). None of these methods are perfect, however. As such, it is advantageous to get a consensus from several different tools before deciding which SNPs to select for further analysis (19).

Although numerous ABCB1 SNPs have been identified in different populations, the functional impact of most nsSNPs in human ABCB1 transporter gene is still unclear. With rapidly increasing in bioinformatics tools and algorithms with better predictive power, computational technologies can aid in predicting nsSNP that are likely to have deleterious effects on ABCB1 protein's structure, its expression, functions or disease susceptibility. In this study, we performed a comprehensive in silico analysis of nsSNPs in coding regions in ABCB1 gene using different structural bioinformatics tools.

## Methodology

### Retrieval of SNPs

The SNP information of human ABCB1 gene was obtained from the National Center for Biotechnology Information (NCBI) SNPs database, (dbSNP) (https://www.ncbi.nlm.nih.gov/SNP/) (accessed Nov.2018). nsSNPs in the coding regions were selected for investigation.

### Prediction of deleterious nsSNPs by different bioinformatics tools

The effects of nsSNPs on ABCB1 protein structure and function were predicted using the following bioinformatics tools: SIFT (http://sift.bii.a-star.edu.sg/) and PolyPhen-2 (http://genetics.bwh.harvard.edu/pph2/index.shtml), were used to predict the deleterious nsSNPs. To increase the accuracy of the in silico techniques for prioritizing deleterious nsSNPs, the nsSNPs found to be deleterious by SIFT & PolyPhen-2 were further analyzed by PANTHER (http://pantherdb.org/), SNAP (https://rostlab.org/services/snap/), PROVEAN (http://provean.jcvi.org) PhD-SNP (http://snps.biofold.org/phd-snp/phd-snp.html) and SNPs & GO (http://snps-and-go.biocomp.unibo.it/snps-and-go/).

Prediction of protein stability changes was analyzed by I-Mutant 3.0 (http://gpcr.biocomp.unibo.it/cgi/predictors/I-Mutant3.0/I-Mutant3.0.cgi). ConSurf (http://consurf.tau.ac.il/) was used to predict the conservation of amino acid positions in a protein molecule. We used NetSurfP server (http://www.cbs.dtu.dk/services/NetSurfP/) to predict the surface accessibility of amino acid residue in protein structure.

Chimera and Project HOPE (http://www.cmbi.ru.nl/hope/) were used to analyze changes in the protein 3D structure due to deleterious nsSNPs. The FASTA format of the protein was obtained from Uniprot at Expassy database.

### Prediction of deleterious nsSNPs by SIFT

SIFT (Sorting Intolerant From Tolerant) tool predicts whether an amino acid substitution at specific position in the protein sequence have a tolerant or deleterious effect on protein function. SIFT automatically perform multiple steps by using NCBI PSI-BLAST: searches for protein sequences homologous to the query protein sequence and obtains multiple alignment of these chosen sequences, then calculates normalized probabilities for all possible substitutions. Substitutions with normalized probabilities less than a chosen cutoff score of 0.05 are predicted to be deleterious and those with scores greater than or equal to the cutoff score are predicted to be tolerated (20–22).

### Prediction of deleterious nsSNPs by SNAP

SNAP (Screening for Non-Acceptable Polymorphisms) is a neural network-based online tool that classify nsSNPs into deleterious (affect protein structure and function) and neutral (no effects) polymorphisms. SNAP provides a reliability index that reflects the level of confidence of a particular prediction (21, 23).

### Prediction of deleterious nsSNPs by PANTHER

PANTHER (Protein Analysis Through Evolutionary Relationships) is a protein family and subfamily database that predicts whether the SNPs will cause a deleterious functional effect on the protein. PANTHER output is subPSEC (substitution position specific evolutionary conservation) score which is negative logarithm of the probability ratio of the wild-type and mutant amino acid at a particular position. The subPSEC scores are continuous values from 0 (neutral) to about 10 (most likely to be deleterious) (24).

### Prediction of damaging nsSNPs by PolyPhen

PolyPhen (Phenotyping Polymorphism) is a web server that automatically predicts whether an amino acid substitution due to coding nsSNPs is damaging (affect protein structure and function) or benign (no effects). PolyPhen performs its prediction based on the 3D protein structure and multiple alignments of homologous sequences, and then calculates Position-Specific Independent Count (PSIC) scores for each of the two variants. The higher the difference of the PSIC scores of the two variants, the higher functional impact a particular amino acid substitution is likely to have. PolyPhen scores are classified as probably damaging (0.95–1), possibly damaging (0.7–0.95) and benign (0.00–0.31) (25–27).

### Predication of deleterious nsSNP by PROVEAN

PROVEAN (Protein Variation Effect Analyzer) predicts the functional impact of not only single amino acid substitutions but also other classes of protein sequence variations including insertions, deletions, and multiple substitutions. PROVEAN collects a set of homology supporting sequences by searching a protein database for related sequences and selects a supporting set from the sequences. Then make a prediction based on a PROVEAN score, with score ≤ -2.5 indicates that the protein variant is deleterious, whereas scores > -2.5 is considered as a neutral variant (28).

### Prediction of Disease related nsSNP by PhD-SNP

PHD-SNP is a support vector machine (SVM) based online classifier that optimized to classify a single point protein mutation into disease-related or neutral polymorphism (29).

### Prediction of Disease Related nsSNP by SNPs & GO

SNPs & GO is a SVM based tool that predicts whether human nsSNPs are disease-associated or neutral based on an accurate method that include Gene Ontology (GO) protein sequence-associated terms (30).

### Prediction of nsSNPs’ Impact on Protein Stability by I-Mutant 3.0

I-Mutant is a SVM based web server that predicts protein stability changes upon single point mutation starting from the protein structure or sequence. I-Mutant provides a free energy change value (DDG) and its direction, where positive value indicates that the mutated protein is of higher stability than the negative value. The DDG value classified into largely unstable (DDG < −0.5 kcal/mol), largely stable (DDG>0.5kcal/mol), or neutral (-0.5e DDGe0.5 kcal/mol) (31, 32).

### ConSurf

ConSurf is a web-server that used phylogenetic of homologous sequences to predict the conservation of amino acid positions in a protein. The conservation scores are nine grades ranging from grade 1 (the most variable positions, turquoise color) to grade 9 (the most conserved positions, maroon color), with grade 5 being the intermediately conserved position (white color) (33).

### In silico biophysical analysis of nsSNPs by NetSurfP

NetSurfP is a neural network web-server that predicts the relative surface accessibility of an amino acid residue in a protein sequence (i.e. predict whether an amino acid is buried or exposed). The reliability of this prediction is reflected by Z-score. NetSurfP also predicts the secondary structure of amino acids in a sequence (34).

### Prediction of protein structure by Project HOPE and Chimera

#### Project HOPE software

HOPE (Have (y)Our Protein Explained) is web server that analyzes the effects of the mutation on 3D structure and function of the protein. HOPE collects information from different data sources including protein’s 3D structure, UniProt database and DAS-servers. Hope processes these data and generates an extensive report that explains and illustrates the effects of the mutation with figures and 3D structural visualization of mutated proteins (35).

#### Chimera software

Chimera software is a program for interactive visualization and analysis of protein 3D structure. Chimera (version 1.8) is available within the Chimera package and available from the Chimera web site (http://www.cgl.ucsf.edu/chimera/) (32, 36). Chimera was used to generate the mutated models of ABCB1 protein 3D model.

## Results

### Retrieval of SNPs

The nsSNPs of ABCB1 gene investigated in this work were retrieved from dbSNP database. It contained a total of 47007 SNPs: 1001 were nsSNPs, 429 were coding synonymous, 693 were in non-coding regions, which comprises of 574 SNPs in 5’ untranslated region (UTR) region and 119 SNPs in 3’ UTR. The rest were in the intron region. The non-synonymous coding SNPs were selected for our investigation.

### Prediction of tolerated and deleterious nsSNPs by SIFT

SIFT analysis predicted that a total of 108 nsSNPs were damaging (score 0.05) and 253 nsSNPs had tolerated effects on the ABCB1 gene.

### Prediction of damaging nsSNPs by PolyPhen-2

According to our Polyphen-2 results, four nsSNPs were predicted as “probably damaging” (rs1128501, rs2235036, rs28381902, rs55852620), and one nsSNP as possibly damaging (rs9282565). To increase the accuracy of predictions, results of SIFT and PolyPhen-2 were joined and SNPs with PolyPhen score> 0.90 and SIFT< 0.05 were selected. Accordingly, three nsSNPs passed both criteria and were classified as deleterious/damaging (Table 1).

**Table 1:** List of nsSNPs that predicted as deleterious by both SIFT and PolyPhen-2.

| dbSNP rs# | Protein residue | SIFT score | SIFT Prediction | PolyPhen Prediction | Polyphen score |
| --- | --- | --- | --- | --- | --- |
| rs28381902 | E566K | 0.00 | deleterious | Probably damaging | 1 |
| rs1128501 | G185V | 0.002 | deleterious | Probably damaging | 1 |
| rs9282565 | A80E | 0.007 | deleterious | Possibly damaging | 0.695 |

### Prediction of deleterious nsSNPs by other bioinformatics tools

PANTHER, SNPs & GO, PhD-SNP, SNAP and PROVEAN confirmed the damaging effect of two nsSNPs (rs28381902 and rs1128501), whereas the nsSNPs rs9282565 found to be deleterious by only PhD-SNP and SNAP (Table 2).

**Table (2):** Prediction of deleterious nsSNPs by different bioinformatics tools.

| dbSNP rs# |  | rs28381902 | rs1128501 | rs9282565 |
| --- | --- | --- | --- | --- |
| Amino acid change |  | E566K | G185V | A80E |
| SNP&GO | Prediction | Disease | Disease | neutral |
|  | RI | 5 | 0 | 1 |
| PhD-SNP | Prediction | Disease | Disease | Disease |
|  | RI | 7 | 2 | 2 |
|  | Probability | 0.429 | 0.524 | 0.751 |
| SNAP | Prediction | Effect | Effect | Effect |
|  | Score | 82 | 57 | 45 |
| PANTHER | Prediction | Disease | Disease | Neutral |
|  | RI | 9 | 2 | 2 |
| PROVEAN | Prediction | Deleterious | Deleterious | Neutral |
|  | Score | -3.514 | -7.513 | -1.417 |

RI, reliability index.

### Prediction of nsSNPs’ Impact on Protein Stability by I-Mutant 3.0

The three nsSNPs were predicted by I-Mutant as significantly decreasing the stability of the protein (E566K, DDG -0.83 Kcal/mol; G185V, DDG -0.73 Kcal/mol; A80E, DDG -0.8 Kcal/mol)

### ConSurf

Analysis of the three deleterious nsSNPs with consurf revealed that all of them were located in highly conserved regions and predicted to have functional and structural impacts on ABCB1 protein (E566K, (score 5, intermediately conserved); G185V, (score 7, highly conserved); A80E, (score 9, highly conserved)).

### In silico biophysical analysis of nsSNPs by NetSurfP

Using NetSurfP, the biophysical stability of the three nsSNPs (E566K, G185V, A80E) was analyzed. As shown in Table 3, a huge drift in the Z-score was not observed for the three nsSNPs. Also, for any of the three, nsSNPs, the class assignment does not change.

**Table 3:**
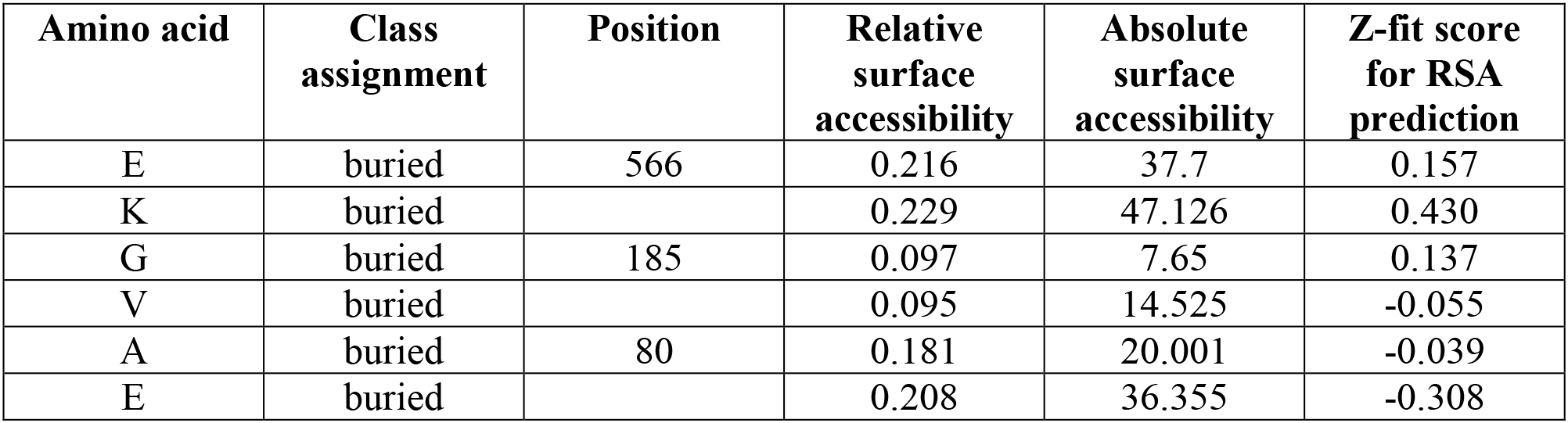
Surface accessibility of wild type and mutant variants in ABCB1.

### Prediction of protein structure by Project HOPE and Chimera

#### Project HOPE software

The 3D analysis of the wild type and mutant protein structures was performed by project HOPE.

#### Mutation (rs28381902) of Glutamic Acid into Lysine at position 566 (E566K)

For this variant, the mutated residue is larger; this might lead to loss of interactions. The wild-type residue was negatively charged, forms a hydrogen bond with Threonine at position 558, and forms a salt bridge with Arginine at position 588. The difference in charge will disturb the ionic interaction made by the original, wild-type residue. The charge of the buried wild-type residue is reversed by this mutation; this can cause repulsion between residues in the protein core and affects protein folding

#### Mutation (rs1128501) of Glycine into Valine at position 185 (G185V)

For this variant, the mutated residue is bigger, this might lead to loss of interactions.it is less hydrophobicthan the wild type. The wild-type residue is a glycine, the most flexible of all residues. This flexibility might be necessary for the protein’s function (37). So, molecular and structural variations between Glycine and Valine can lead to different conformational changes thus it mediates the distortion and structural de-stability due to less flexibility.

#### Mutation (rs9282565) of Alanine into Glutamic Acid at position 80 (A80E)

The wild-type residue charge was neutral; the mutant residue charge is negative. Introduction of a charge in a buried residue can lead to protein folding problems. The wild-type residue is more hydrophobic than the mutant residue. The mutation may cause loss of hydrophobic interactions in the core of the protein.

### Chimera Software

Chimera was used to visualize changes in the protein 3D structure due to deleterious nsSNPs.

**Figure 1:**
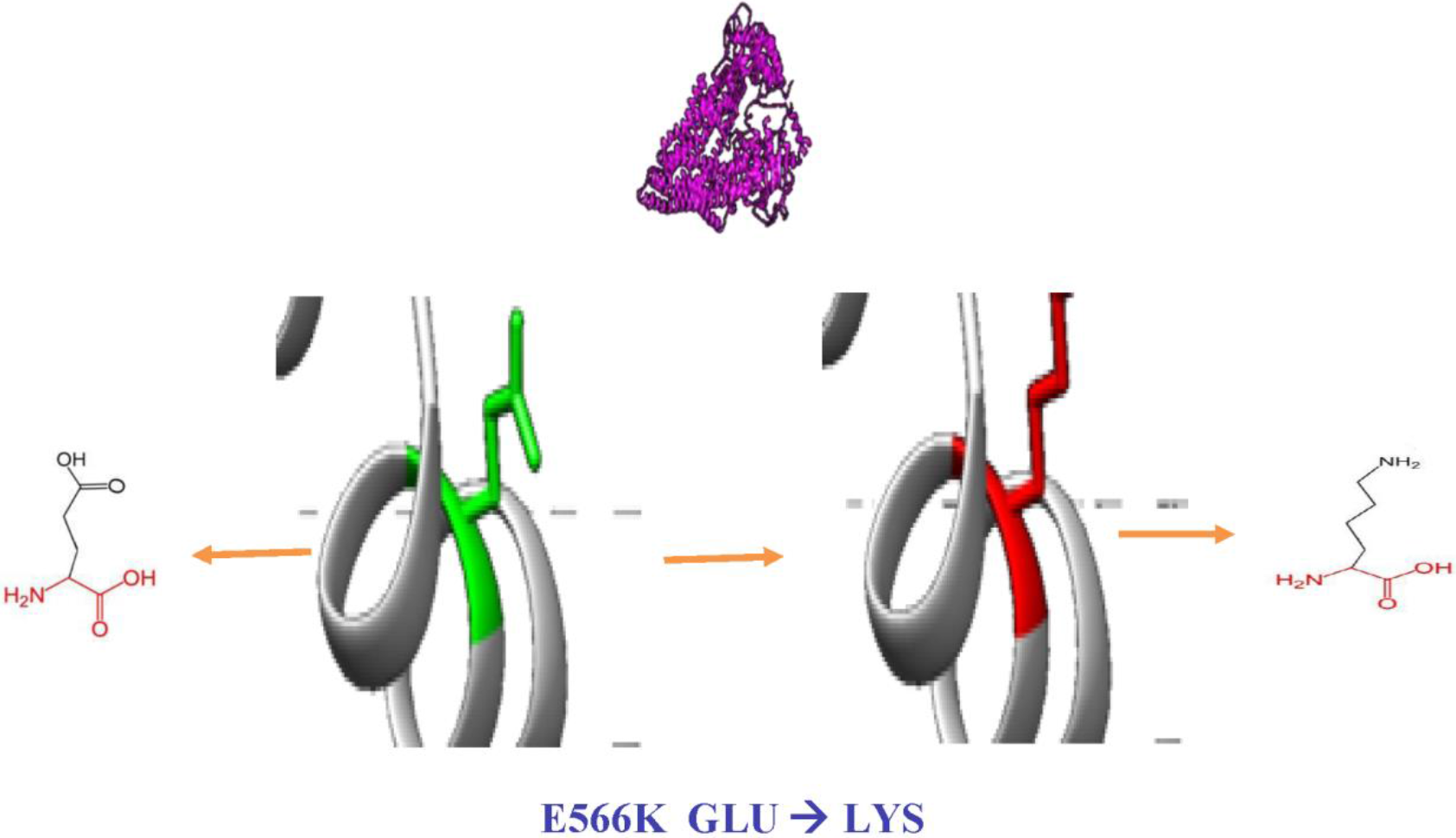
nsSNP rs28381902 change the amino acid Glutamic Acid (green color) into Lysine (red color) at position 566.

**Figure 2:**
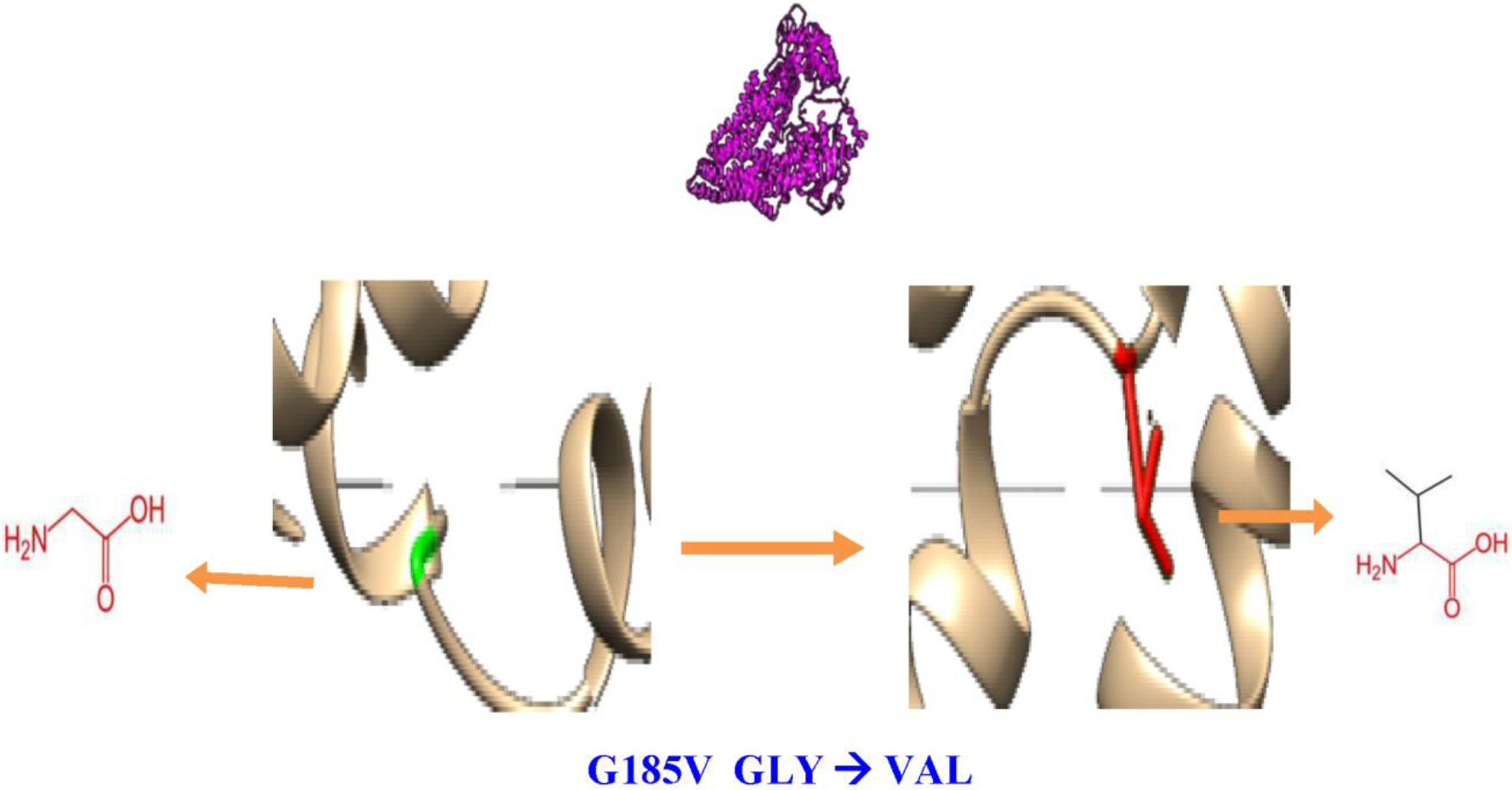
nsSNP rs1128501 change the amino acid Glycine (green color) into Valine (red color) at position 185.

**Figure 3:**
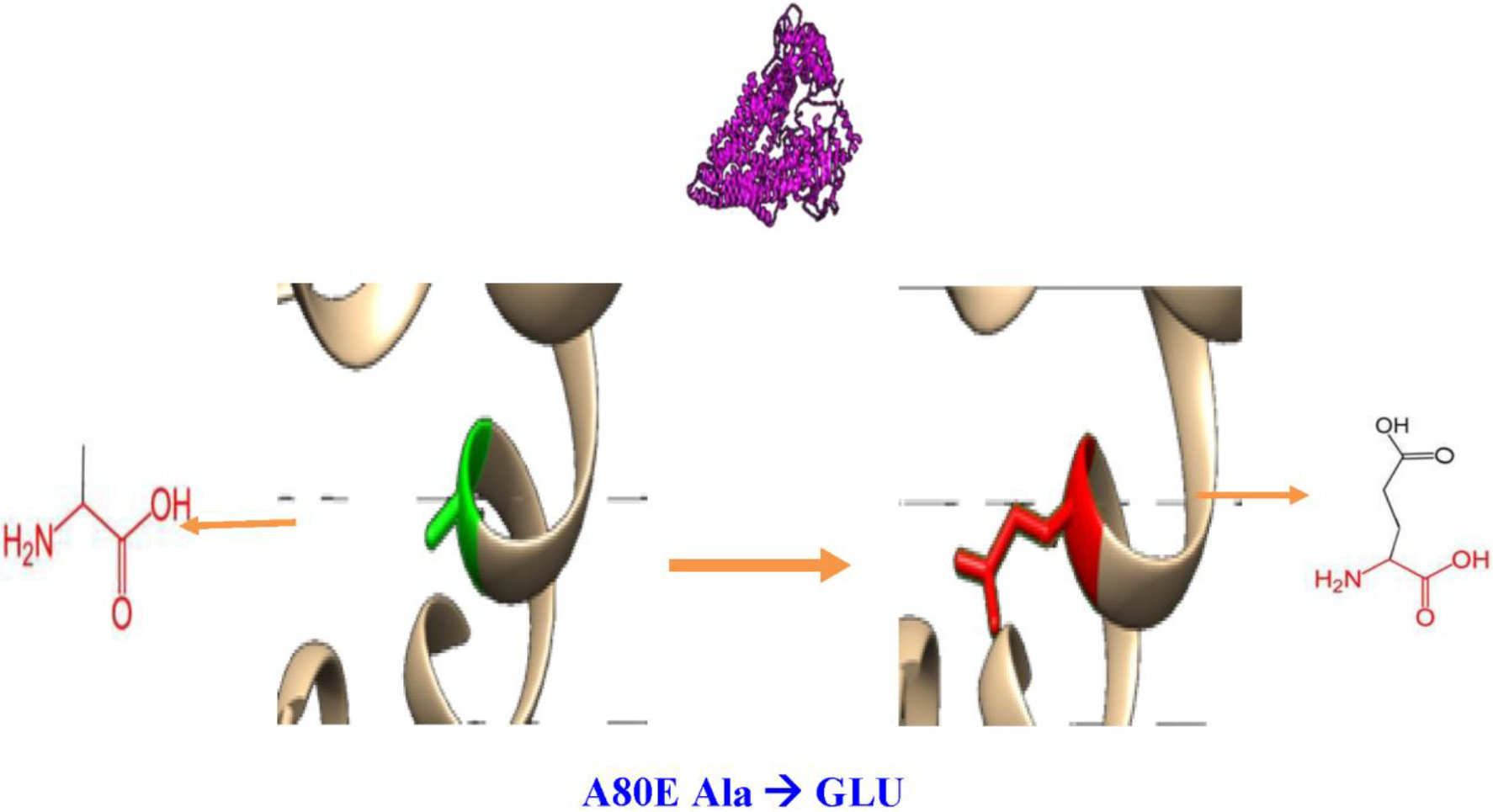
nsSNP rs9282565 change the amino acid Alanine (green color) into Glutamic Acid (red color) at position 80.

## Discussion

ABCB1 plays a key role in transport of exogenous and endogenous molecules across cell membrane (1–7). Increasing evidence has suggested that functional SNPs of ABCB1 gene may contribute to many diseases (2, 5–7, 10) as well as response to many drugs (2, 5–7, 11–18, 38–40). So, an effort was made to identify SNPs that can modify the structure, function and expression of the ABCB1 gene. Therefore, discriminating deleterious SNPs with potential effects on disease susceptibility from tolerated variants is a major challenge. Therefore, a comprehensive study that systematically analyzes the effects of such SNPs can cost-effectively prioritized SNPs for further analysis.

Previous studies on polymorphisms screening using in silico analysis helped in predicting the functional nsSNPs associated with genes. The studied genes include MC1R (41). Human TNF alpha (42), human MBL-2 (43), ABCB1 and ABCC1 in *Bos Taurus*, human proliferating cell nuclear antigen (44), oestrogen receptor (45) and TLRs signaling pathway (33).

Most of the SNPs of the *ABCB1* gene have not been studied yet. In this study, a computational approach was undertaken to systematically analyze nsSNPs to predict the deleterious mutations. The strengths and weaknesses and the differences in the predictions generated by different bioinformatics programs indicate the need for a combined analysis to enhance the accuracy of effect predictions (19, 46). As a general rule, at least 4 or 5 of these tools should be run to gain a consensus on the effect of the SNP (19). So, the nsSNPs situated in the ABCB1 gene were evaluated by seven programs that use different methods to predict the damaging nsSNPs. Those include tools that make predictions based solely on the sequence of a protein (SIFT, PROVEAN, PANTHER, PhD SNP, SNP&GO) and tools that incorporate structural information when making predictions on the functional effects of mutations (PolyPhen, SNAP).

SIFT and PolyPhen were reported to have better performance in identifying deleterious nsSNPs (47). The accuracy of SIFT and PolyPhen 2.0 was further validated by (Hicks, et al., 2011) (48), which makes these tools more applicable for the prediction. Based on these in silico studies, this study has tested the nsSNPs of ABCB1 gene using SIFT and PolyPhen. The nsSNPs with double positive results were considered as the most damaging, which were then analyzed suing I-Mutant, PROVEAN, SNP&GO, PHD-SNP and PANTHER for the screening of functional mutation in ABCB1 gene. By comparing the scores of the seven methods, three nsSNPs with positions E566K, G185V and A80E were found to be highly deleterious with agreement between at least four tools. This result suggests that these variants may be considered to be the most likely damaging or deleterious nsSNPs.

The analysis of protein stability change allows confirmation of this finding. In fact, using I-mutant program, the three nsSNPs were found to induce a decrease in energy in comparison with the native structure. Netsurf and Consurf were used to assess the surface accessibility and the conservation of the protein, respectively. HOPE showed changes in protein secondary structure, and chimera was used to visualize the 3D structure of the protein.

In conclusion, available databases such as NCBI and dbSNP, along with in silico prediction programs were surveyed and compared to assess the effects of deleterious nsSNPs on the ABCB1 protein functions. These programs are based on evolutionary, structural and computational methods. Hence, implementation of different algorithms often serves as powerful tools for prioritizing candidate functional nsSNPs for further studies.

This study predicted three nsSNPs in the ABCB1 gene as the most damaging. These nsSNPs, to the best of our knowledge, have not yet been investigated and therefore may be considered as candidates for association with diseases and drug response.

## Supporting information

Supplemental Table 2: Pholyphen result

Supplemental Table 1: SIFT result

